# Bridging multiple scales in the human brain using computational modelling

**DOI:** 10.1101/085548

**Authors:** Michael Schirner, Anthony Randal McIntosh, Viktor K. Jirsa, Gustavo Deco, Petra Ritter

## Abstract

Brain dynamics span multiple spatial and temporal scales, from fast spiking neurons to slow fluctuations over distributed areas. No single experimental method links data across scales. Here, we bridge this gap using The Virtual Brain connectome-based modelling platform to integrate multimodal data with biophysical models and support neurophysiological inference. Simulated cell populations were linked with subject-specific white-matter connectivity estimates and driven by electroencephalography-derived electric source activity. The models were fit to subject-specific resting-state functional magnetic resonance imaging data, and overfitting was excluded using 5-fold cross-validation. Further evaluation of the models show how balancing excitation with feedback inhibition generates an inverse relationship between α-rhythms and population firing on a faster time scale and resting-state network oscillations on a slower time scale. Lastly, large-scale interactions in the model lead to the emergence of scale-free power-law spectra. Our novel findings underscore the integrative role for computational modelling to complement empirical studies.

---

Empirical approaches to characterizing the neural mechanisms that govern brain dynamics often rely on the simultaneous use of different acquisition modalities. These data can be merged using statistical models, but the inferences are constrained by the scales of measurement, rendering a mechanistic understanding of multiscale dependencies elusive. Mathematical modelling has played a central role for interpreting multiscale data in many areas of science, including physics, chemistry, and biology. Computational modelling accelerates the identification of principles that bridge spatiotemporal scales serving as the quantitative link between experimental results^1, 2^.

Here, we use the open-source connectome-based modelling platform The Virtual Brain^3–7^ (thevirtualbrain.org) to develop large-scale brain network models. The innovation in this paper is to model individual brains using empirical neuroimaging data to constrain large-scale dynamics. Person-specific white-matter connectivity sets the framework for individual dynamical systems models that are driven with electroencephalography (EEG) source imaging activity derived from the same persons. The source activity serves as biologically-based approximation of synaptic input currents^8–10^ and replaces the more common use of noise-driven dynamics (Fig. 1).

The modelling effort reported here addresses three frequently reported empirical phenomena: *(i)* negative correlations between resting state functional magnetic resonance imaging (fMRI) and EEG α-power fluctuations observed by simultaneous EEG-fMRI^11–13^; *(ii)* α-rhythm based gating by inhibition^14, 15^; and *(iii)* scale-freeness of fMRI and EEG signals^16–18^ as illustrated in **Supplementary Video 1**.

A recent focus of neuroscience has been on intrinsic neural dynamics and their relation to brain structure and functional capacity^19–21^. Much of this was motivated by observations from fMRI studies of the so-called “Resting-State Networks” (RSN), which showed coherent spatiotemporal dynamics in the absence of an explicit task^22–24^. Support for a neuronal basis comes from concurrent fMRI and intracortical recordings^25, 26^, simultaneous EEG-fMRI^11–13, 27, 28^ and magnetoencephalography^29, 30^ studies that independently detected the spatial patterns of RSNs. Several studies report negative correlations between cortical fMRI blood oxygen level–dependent (BOLD) contrast signals and EEG-derived α-rhythm power^11, 12, 31, 32^. EEG α-rhythms have traditionally been interpreted as a sign of cortical “idling”^33^. This notion is now extended by a growing body of research that emphasizes their role for information processing^34^. Importantly, attention, perceptual awareness and cognitive performance are rhythmically modulated by α-power and phase^35–38^. The roles of α-rhythms for mediating top-down control, timing of oscillations and directing attention by blocking task-irrelevant pathways are central to prevailing hypotheses termed ‘gating by inhibition’ and ‘pulsed inhibition’^14, 15^. Intracellular recordings showed that inhibition is inseparable from excitatory events, resulting in ongoing excitation-inhibition balance (E/I balance)^39, 40^. Finally, despite widespread interest in criticality and its ubiquity in nature^18^ the origin of power-law scaling observed in both fMRI and EEG data is unclear^16, 17^. A key point of consideration across the three phenomena we describe in the context of the present paper is that the computational model was not explicitly constructed to address each of them. Rather we used an empirically constrained connectome-based brain network model to provide plausible multiscale explanations that link these phenomena and uncover underlying mechanisms.

## Prediction of individual fMRI time series

**Figure 1.**
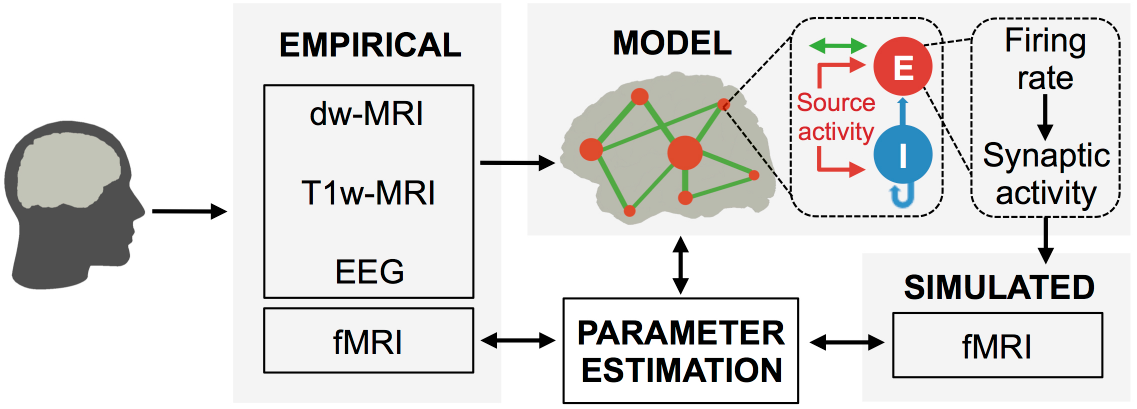
**Hybrid modelling framework.** Person-specific hybrid brain network models are networks of local neural population models that are driven by empirical activity, rather than noise. Population models represent the activity of individual brain regions. These local models are globally coupled by structural connectomes extracted from diffusion-weighted MRI (dw-MRI) using brain parcellations obtained from T1-weighted MRI (T1w-MRI). Population models are injected with region-wise EEG source activity approximating locally generated synaptic input currents^9, 10^. This enables person-specific simulations. Simulated whole-brain fMRI time series are fitted with empirical fMRI time series that were acquired simultaneously with EEG. 5-fold cross-validation was performed to guard against overfitting. The underlying simulated neural activity (firing rates and synaptic activity) is analysed to reveal links between mesoscopic dynamics and neuroimaging signals. Excitatory postsynaptic currents (EPSCs) tend to be strongly correlated with the electric fields in their vicinity, while inhibitory postsynaptic currents (IPSCs) are anticorrelated with EPSCs^8, 39, 40^. Here, EPSCs are derived from EEG and IPSCs result from modelled inhibitory population activity. Synaptic gating (the fraction of open channels of neural populations) was transformed to region-wise fMRI signals using the Balloon-Windkessel forward model^41^. In contrast to noise-driven models that only capture stationary features, hybrid models as used here are able to predict the ongoing temporal dynamics of fMRI time series.

Using exhaustive searches, we fitted parameters for brain network models of 15 human adult subjects, who all had T1-weighted, diffusion-weighted and simultaneous resting state EEG-fMRI data (Fig. 1). For each brain network model, three parameters were varied to maximize the fit between empirical and simulated fMRI: the scaling of excitatory white-matter coupling and the strengths of the inputs injected into excitatory and inhibitory populations. Besides tuning these three global parameters to maximize fMRI fit, we also tuned all local inhibitory coupling strengths in order to obtain biologically plausible firing rates in excitatory populations. For this second form of tuning, termed feedback inhibition control (FIC), average population firing rates were the sole optimization criterion, without any consideration of prediction quality, which was only dependent on the three global parameters (see Methods). FIC is required to compensate for excess or lack of excitation resulting from the large variability in white-matter coupling strengths obtained by MRI tractography, which is a prerequisite to obtain plausible ranges of population activity that is relevant for some of the following results (Figs. 4 and 5). Prediction quality was measured as the average correlation coefficient between all simulated and empirical region-wise fMRI time series of a complete cortical parcellation over 20.7 minutes length (TR = 1.94s, 640 data points) thereby quantifying the ability of the model to predict the activity of 68 parcellated cortical regions. Accounting for the large-scale nature of fMRI resting-state networks, the chosen parcellation size provides a parsimonious trade-off between model complexity and the desired level of explanation. What this parcellation may lack in spatial detail, it gains in providing a full-brain coverage that can reliably reproduce ubiquitous large-scale features of empirical data, which we further present below. To exclude overfitting and limited generalizability, we performed 5-fold cross-validation. For each subset (fold), the fMRI data were randomly divided into an 80% sample as training segment and a 20% sample as testing segment. Furthermore, despite that large range of possible parameters, the search converged to a global maximum (Supplementary Fig. 1). Therefore, we ensured that when the model has been fit to a subset of empirical data, that it was able to generalize to new or unseen data. In contrast to model selection approaches, where the predictive power of different models and their complexity are compared against each other, we here use only a single type of model.

The hybrid model simulation results were compared with three control scenarios: *(i)* a model where local dynamics were driven by noise, *(ii)* a variant of the hybrid model that used random permutations of the region-wise EEG source activity time series and *(iii)* a statistical model where ongoing α-band power fluctuation of EEG source activity was convoluted with the canonical hemodynamic response function (henceforth called α-regressor). The first two controls are brain network models; the third is inspired by traditional analyses of empirical EEG-fMRI data. The combination of 5-fold cross validation and comparison against these control scenarios further reinforces the explanatory utility of the hybrid computational model.

**Figure 2.**
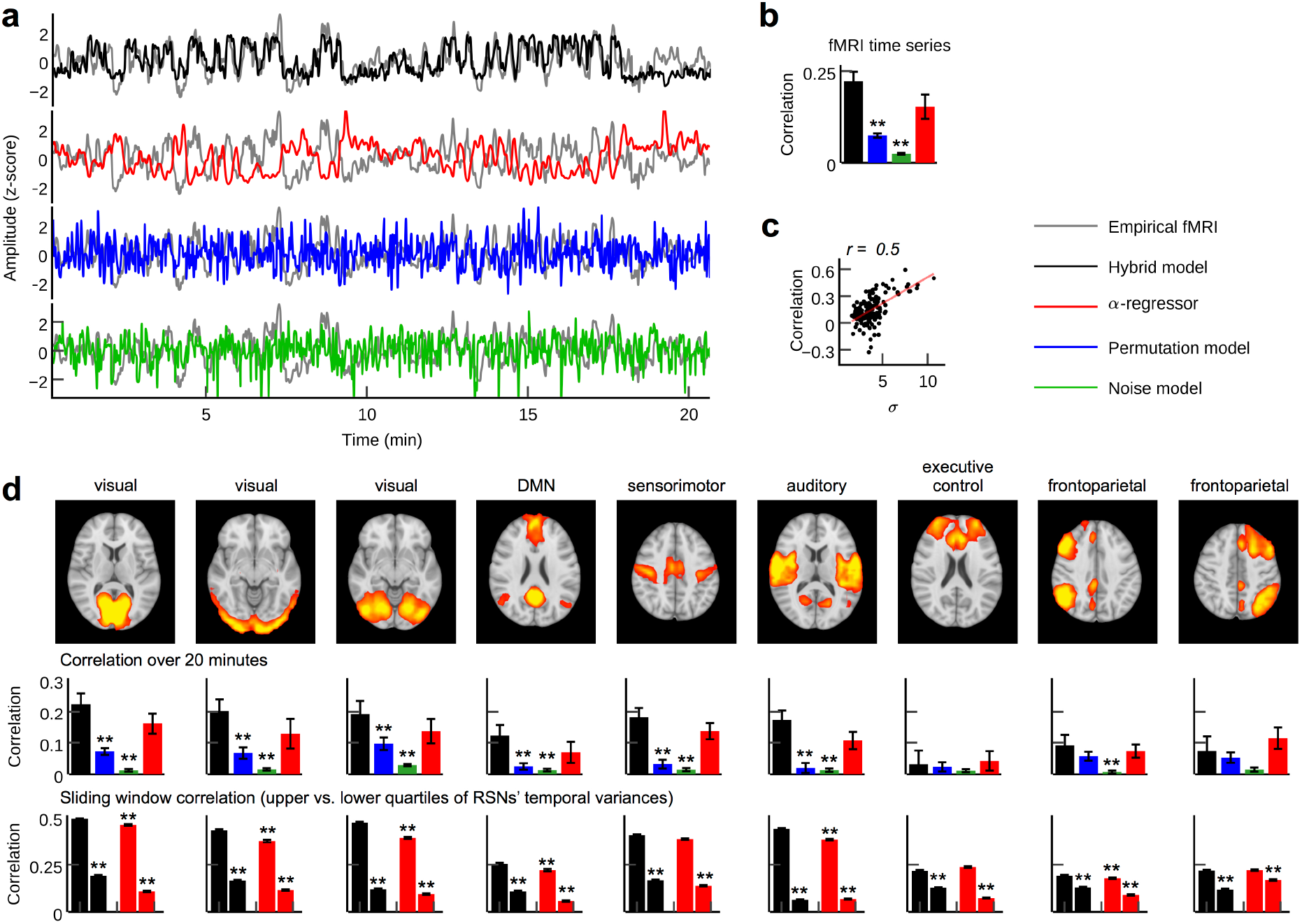
Person-specific fMRI time series prediction. **a**, Example time series of the hybrid model and the three control scenarios. The hybrid model shows good correspondence with empirical fMRI waveforms compared to control network models. The α-regressor correlates comparably strong, but negatively, with empirical fMRI. **b**, Mean values of correlation coefficients between all simulated and empirical region time series (∼20 minutes) for all subjects (errorbars indicate standard errors of the mean; values for the α-regressor were inverted for illustration purposes). Time series predictions from the hybrid model yield significantly higher correlations than control network models. **c**, Standard deviation (s.d.) of RSN time courses correlates with fMRI time series prediction quality. Dots plot the s.d. of each of the nine RSN time courses in each of the 15 subjects versus the prediction quality of fMRI time series for all regions underlying the respective RSN. Prediction quality varies over regions and subjects; the hybrid model predicts an fMRI region time series better when its corresponding RSN time course has a high s.d., i.e., when RSN activity dominates the fMRI signal of that region. **d**, Prediction of the temporal dynamics of RSNs. Upper row: spatial activation patterns of group-ICA RSNs. Middle row: mean correlation coefficients between RSN temporal modes and hybrid model simulation results, respectively α-regressor. Lower row: mean sliding window (length = 100 fMRI scans = 194 s; one fMRI scan step width) correlations for the upper and lower quartiles of window-wise RSN temporal mode s.d. Left and right bars show the average correlations for the upper and lower quartiles of window-wise s.d. for the hybrid model and the α-regressor. Prediction quality of RSN temporal activity is significantly improved during epochs of high RSN activation. Asterisks indicate significant differences compared to the source activity model (*p < .05, **p < .01, two-sample t-test).

Visual inspection of example time series showed good reproduction of characteristic slow RSN oscillations by the hybrid model and the α-regressor (albeit inverted for the latter), but poor reproduction of temporal dynamics in the case of noise and random permutations models (Fig. 2a). To quantify prediction quality, we computed the mean over all correlation coefficients between simulated and empirical fMRI time series for each parameter set and each of the four setups (i.e., hybrid model and the three control setups). Predictions from the hybrid model correlated significantly better with empirical fMRI time series than predictions from the two random models (Fig. 2b). Five-fold cross-validation showed no significant difference of prediction quality between training and validation data sets *(p = .97, two-sample t-test)* and between validation data sets and prediction quality for the full time series *(p = .34, two-sample t-test)*. Predictions from the α-regressor were slightly less accurate (note that as α-power is inversely correlated with fMRI activity, correlation coefficients were inverted for comparison).

We performed a group-level spatial independent component analysis (ICA) on the empirical fMRI data to compare simulation results with RSN activity (Fig. 2d). To account for temporal variation of prediction quality, we computed sliding-window correlations (100 fMRI scans = 194 s window length; one fMRI scan step width). Average prediction quality was largest for the hybrid model for eight out of nine RSNs (Fig. 2d). As with region-wise fMRI, correlation coefficients of the hybrid model and the α-regressor were comparably good, while the control network models performed poorly. We found that prediction quality varied over time, regions and subjects. Window-wise prediction quality was highly correlated with the standard deviation of RSN temporal modes (Fig. 2c). That is, the higher the temporal activation of a RSN (in a given subject, region and time window) the better the prediction of its activation time series. As a consequence, epochs in the upper quartile of RSN standard deviations were significantly better predicted than epochs in the lower quartile (Fig. 2d). Similarly, the strength of functional networks (measured by average FC between all region-pairs) correlated with prediction quality (r=0.5 ± 0.32, mean ± s.d.), indicating that epochs of high functional network activity were better predicted than epochs that are characterized by uncorrelated activity.

**Figure 3.**
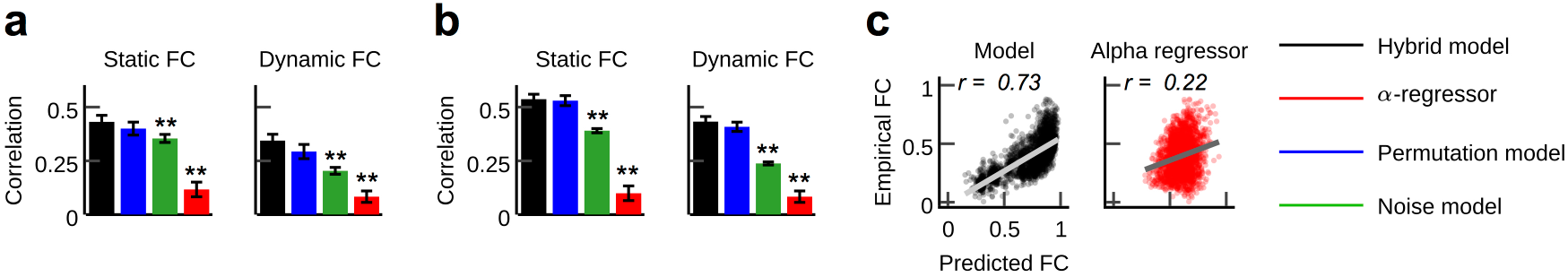
Empirical FC prediction. In contrast to the α-regressor, the hybrid model concurrently predicts empirical time series dynamics (Fig. 2, 4-6) and FC. **a, b**, Mean correlation coefficients between predicted and empirical FC for the three model types and the α-regressor (errorbars indicate standard errors). FC was computed for long (static FC; computed over ∼20 min) and short epochs (dynamic FC; average sliding window correlation; 100 fMRI scans window length; one fMRI scan step width). The hybrid model yields significantly higher individual FC correlations than noise model and α-regressor. Results were compared for the parameter set that generated the best fMRI time series prediction (**a**) and the parameter set that yielded the best FC predictions for each subject (**b**). **c**, Comparison of average FC of empirical data and hybrid model results, respectively α-regressor results. Dots show pairwise FC values for all region-pairs for the full length of the time series averaged over all subjects. Asterisks indicate significant differences compared to the hybrid model (*p < .05, **p < .01, two-sample t-test).

In contrast to time series prediction, the α-regressor performed poorly for estimating FC. Compared to the α-regressor, all three model-based approaches provided significantly better predictions of subject-specific long-term FC (computed over ∼20 min between all 68 ROIs) and functional connectivity dynamics (Fig. 3). Furthermore, FC matrices obtained from hybrid model simulations correlated significantly higher with empirical activity than predictions obtained from the noise-driven model (Fig. 3a, b). Interestingly, correlations for hybrid and random permutation models were effectively the same, likely because the large-scale network dynamics that drive the emergence of FC would be relatively preserved when permuting injected activity. Prediction of group-average FC (all pairwise FC values averaged over all subjects) was higher for the hybrid model compared to the α-regressor (Fig. 3c).

## E/I balance generates α-phase related firing

After fitting the 15 person-specific hybrid models, we analysed the local population activity underlying predicted fMRI time series to infer the neurodynamic mechanisms. Simulated dynamics result from fitting three global model parameters: the scaling of global white-matter coupling and the strengths of inputs injected into excitatory and inhibitory populations. The optimized hybrid model reproduced the inverse relationship between α-phase and firing rates observed in invasive recordings^42^ (Fig. 4a). To investigate these fast-acting dynamics related to α-phase, we computed grand average waveforms of modelled synaptic inputs, population firing rates and synaptic gating, all time-locked to the zero-crossings of α-cycles (Fig. 4b). Resulting waveforms illustrate the relation of α-oscillations to population activity and how the ongoing balancing of rhythmic excitatory and inhibitory inputs generated pulsed inhibition.

**Figure 4.**
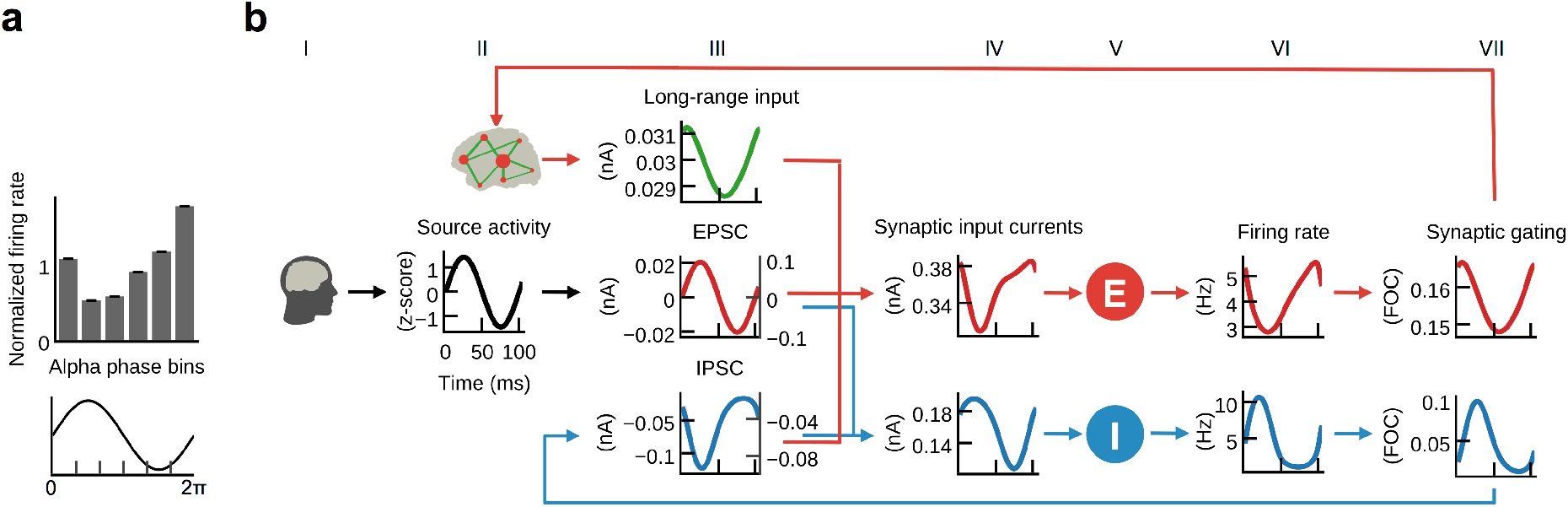
Excitation-inhibition balance underlies the empirically observed inverse relationship between neuronal firing and α-oscillations^42^. Hybrid models that were fitted to person-specific fMRI time series reproduced the invasively observed rhythmic inhibition of firing relative to α-cycle phase (pulsed inhibition) on a faster time scale. **a**, Population firing rates were highest during the trough and lowest during the peak of α-cycles. Firing rates were divided into six bins according to α-cycle segments and normalized relative to the mean firing rate of each cycle. The observed pattern resembles empirical findings^42^. **b**, Grand average waveforms (shaded areas indicate standard errors of the mean) of population inputs and outputs were computed time locked to α-cycles of EEG source activity (black, column II). In optimized models, inhibitory population inputs (blue, column III) were dominated by EPSCs (red); left axes denote currents going into excitatory populations, right axes denote currents going into inhibitory populations. Consequently, firing rates and synaptic gating of inhibitory populations (blue) in columns VI and VII closely followed source activity shape (FOC (column VII) = fraction of open channels). As a consequence, IPSCs resulting from feedback inhibition (blue, column III) showed an inverted pattern; their amplitude at excitatory populations was about three times larger than EPSCs, also reproducing empirical observations^8, 43^. Accordingly, excitatory population input, firing rates and synaptic gating (red; columns IV, VI, VII) were inverted relative to the α-cycle.

As noted above, we injected region-wise EEG source activity into corresponding network nodes in order to simulate biologically realistic EPSCs (Fig. 1). In hybrid models, which were optimized for highest empirical fMRI time series fit and biological plausibility of average firing rates, EPSCs dominated inputs to inhibitory populations. Consequently, the sum of synaptic input currents to inhibitory populations closely followed the shape of EPSCs. As a result of the monotonic relationship between input currents and output firing rates of populations (defined by Eqs. 3 and 4, see Methods), the waveform of inhibitory firing rates and synaptic gating also closely followed injected EPSCs. As increasing input to inhibitory populations leads to increasing inhibitory effect and vice versa, resulting feedback inhibition (i.e. IPSC) waveforms were inverted to injected EPSCs; that is, excitation and inhibition were balanced during each cycle, which is in accordance with published electrophysiology results^8, 40^. Consequently, IPSCs peaked during the trough of the α-phase and were lowest during the peak of the α-phase. The previously fitted synaptic coupling strength parameters determined the relative contribution of excitation and inhibition. Fitting the models to fMRI activity resulted in a biologically plausible ratio^8, 43^ of EPSCs to IPSCs, with IPSC amplitudes being about three times larger than EPSC amplitudes (Fig. 4b). As per other empirical observations^8, 43^, we ensured the EPSC amplitudes at excitatory populations are smaller than IPSC amplitudes by constraining parameters such that the standard deviation of injected source activity was larger for inhibitory populations, i.e., 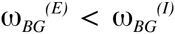. Specifically, we scanned 12 different ratios of both parameters 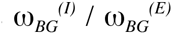 with values between 5 and 200. Because IPSCs have dominated excitatory population inputs, firing rates of excitatory populations showed a similar shape as feedback inhibition currents, i.e., they peaked during the trough of the α-cycle and fell to their minimum during the peak of the α-cycle, reproducing their empirical relationship^42^.

In summary, the mesoscopic activity underlying fMRI predictions showed a rhythmic modulation of firing rates on the fast time scale of individual α-cycles in accordance with empirical observations^42^. Model activity revealed that periodic microstates of excitation and inhibition resulted from the ongoing balancing of EPSCs by feedback inhibition currents.

## ‘Pulsed inhibition’ generates fMRI oscillations

We next focused on the interaction between α-power fluctuations and population dynamics to identify the mechanism underlying the inverse relationship between α-power and fMRI^12^. To isolate the effects of α-waves and separate it from other EEG rhythms, we replaced the EEG-source activity in the optimized model for each subject with artificial α-activity consisting of a 10 Hz sine wave that contained a single brief high power burst in its centre (using the single parameter set for which the highest average prediction quality over all subjects was obtained, Supplementary Fig. 1) and computed grand average waveforms over all resulting region time series (Fig. 5a). As for fMRI prediction, local inhibition strengths for each node were tuned using FIC to obtain biologically realistic firing rates.

**Figure 5.**
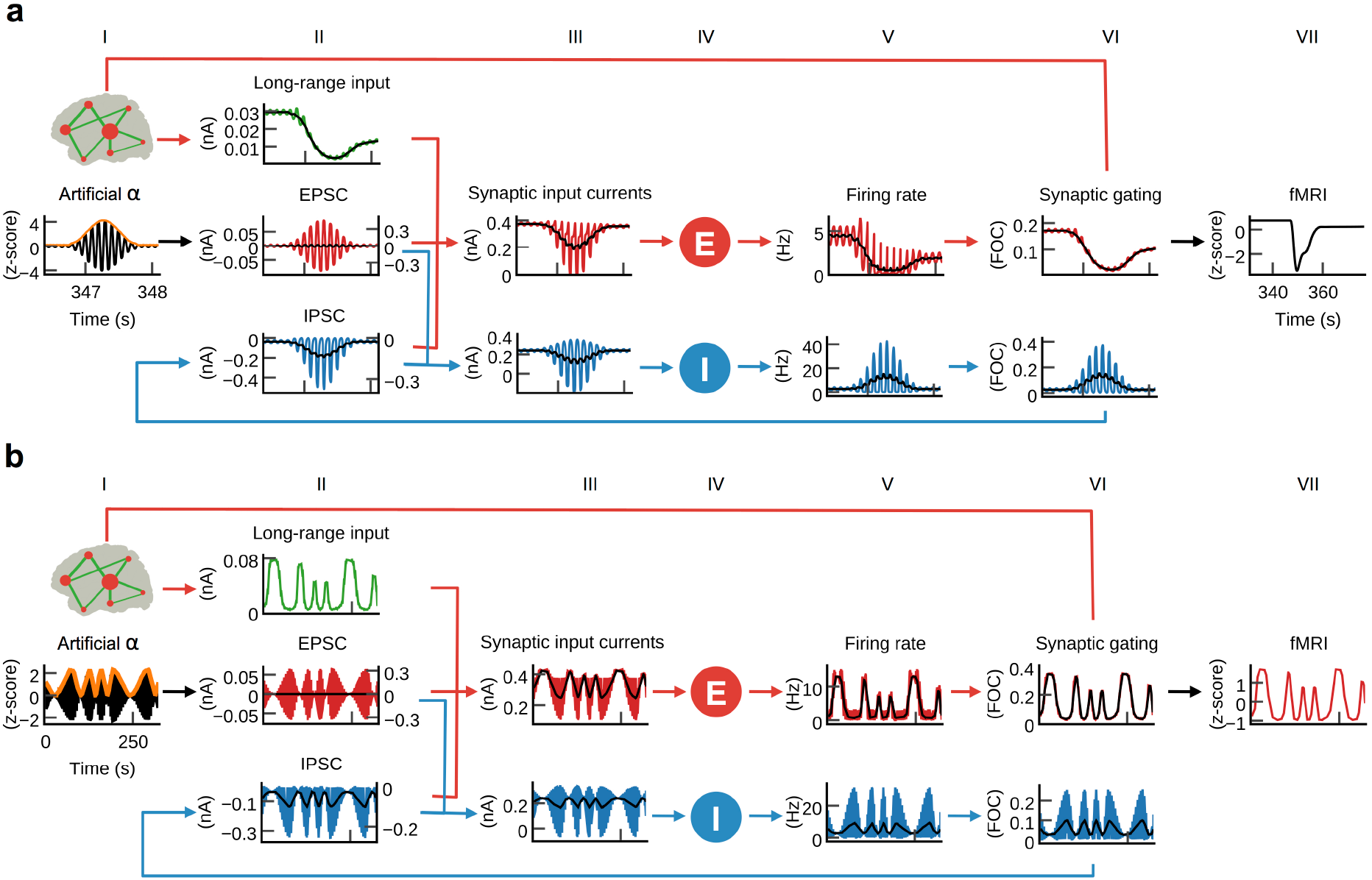
α-power fluctuations generate alternating states of excitation and inhibition that lead to fMRI oscillations. **a**, To isolate the mechanism underlying α-power-related inhibition, subject-specific models were injected with artificial α-activity. Each region received ten minutes of 10 Hz sine oscillations that contained a single brief high power burst (first column; black: centre part of injected artificial α-activity, orange: α-power). While positive deflections of the α-wave generated positive deflections of ongoing firing rates of inhibitory populations, large negative deflections were bounded by the physiological constraint of 0 Hz (blue, fifth column; moving average in black). That is, inhibitory populations rectified α-oscillations such that only the positive half of the wave remained. As a result, average per-cycle feedback inhibition increased for increasing α-power, and the average firing rates, synaptic gating and ultimately fMRI signals (red; fifth, sixth and seventh column) decreased. **b**, To identify the relationship between fMRI and α-rhythms we injected models with 10 Hz sine waves where ongoing power was modulated similar to empirical α-rhythms (0.01 – 0.03 Hz). Similarly to **a**, but for a longer time frame, inhibitory populations rectified negative deflections, which introduced the α-power modulation as a new frequency component into firing rates and fMRI time series.

We analysed waveforms of model state variables and found, as expected, that average per-cycle firing rates and fMRI activity decreased in response to the α-burst (Fig. 5a). Notably, this behaviour emerged despite the fact that injected activity was centred at zero, i.e., positive and negative modulations of input currents were balanced. The reason for the observed asymmetric response to increasing power levels originated from inhibitory population dynamics: while positive deflections of α-cycles generated large peaks in ongoing firing rates of inhibitory populations, negative deflections were bounded by 0 Hz (Fig. 5a). Because of this rectification of high-amplitude negative half-cycles, average per-cycle firing rates of inhibitory populations increased with increasing α-power. As a result, also feedback inhibition had increased for increasing α-power, which in turn led to increased inhibition of excitatory populations, decreased average firing rates, synaptic gating variables and ultimately fMRI amplitudes.

We next analysed the relationship between fMRI oscillations and α-power. We generated artificial α-activity consisting of a 10 Hz sine wave that was amplitude modulated by slow oscillations (cycle frequencies between 0.01 and 0.03 Hz) and injected it into the models of all subjects using the same parameters as for the single α-burst (Fig. 5b). As in the previous experiment, inhibitory populations filtered negative α-deflections during epochs of increased power. This half-wave rectification led to increased average per-cycle firing rates in proportion to signal power, which introduced a new slow frequency component into the resulting time series. The activity of inhibitory populations can be compared to envelope detection used in radio communication for AM signal demodulation. The new frequency component introduced by half-wave rectification of α-activity modulated feedback inhibition, which in turn modulated excitatory population firing rates. Interestingly, the resulting oscillation of firing rates was propagated to synaptic dynamics where the large time constant of NMDAergic synaptic gating (*τ_NMDA_* = 100 ms vs. *τ_GABA_* = 10 ms) led to an attenuation of higher frequencies. The low-pass filtering property of the hemodynamic response additionally attenuated higher frequencies such that in fMRI signals only the slow frequency components remained. To restate: ongoing α-power fluctuations create a slow phase modulation of firing rates and synaptic activity; the low-pass filtering properties of slow synaptic gating and hemodynamic responses attenuate higher frequencies such that only slow oscillations remain in fMRI signals.

In summary, we found that increased α-power led to an increased activation of inhibitory populations. Resulting feedback inhibition generates fMRI oscillations, which can explain the empirically observed anticorrelation between α-power and fMRI.

## Global coupling controls scale-freeness

Empirical fMRI power spectra follow a power-law distribution *P α f^β^*, where *P* is power, *f* is frequency and *β* the power-law exponent, which is an indicator of criticality and scale-free dynamics^44^. In accordance with systematic analyses^44^, average power-law spectra of our fMRI data obeyed power-law distributions with exponents *β_emp_* = −0.97 (*β_emp_* = −0.82 when the frequency range was limited to frequencies < 0.1 Hz as in Ref [44]) for empirical and *β_sim_* = − 0.76 for simulated data (Fig. 6a and Supplementary Fig. 2); time series were tested for scale invariance using rigorous model selection criteria (see Methods). Our previous results (Fig. 5) associated resting-state fMRI oscillations with EEG by identifying a neural mechanism that transforms instantaneous EEG source power fluctuations into fMRI oscillations. Surprisingly, however, the spectrum of EEG source power had a considerably smaller negative exponent than fMRI (*β_α-band_* = −0.56 for α-power and *β_wide-band_* = −0.53 for wide-band power). Comparison of power spectra indicated that power-law fits of simulated fMRI spectra had a higher negative exponent than power-law fits of instantaneous source power spectra because the power of slower oscillations increased relative to the power of faster oscillations (Fig. 6a and Supplementary Fig. 2). Model dynamics transform input activity such that the amplitude of output oscillations increased inversely proportional to their frequency. The effect is visible in Fig. 5b, where fMRI, synaptic and firing rate amplitudes of slow oscillations were larger than amplitudes of fast oscillations, despite equally large amplitudes of input α-power oscillations. In comparison, simulation results obtained for deactivated large-scale coupling, but an otherwise identical setup, did not show this frequency-dependent amplification (Fig. 6b). Without large-scale coupling the power-law exponent of simulated fMRI (*β_sim_Gzero_* = − 0.53) was close to the exponent of injected EEG source activity power (*β_wide-band_* = −0.53). We studied the link between large-scale excitation, local inhibition and scale-free dynamics by analysing simulation results for different settings of global coupling and local feedback inhibition. To isolate the effect of global coupling and exclude that this frequency-dependent amplification is due to feedback inhibition control (see Methods) and to disentangle the effects of global excitation and local inhibition, we used a single weight for all inhibitory couplings and obtained a consistent, but more drastic effect: when global coupling and FIC were both deactivated, simulated fMRI showed strongly decreased scale invariance (the scaling exponent *α = 0.56* obtained by detrended fluctuation analysis was close to *α = 0.5*, which indicates uncorrelated white noise), which demonstrated the crucial role of global coupling for the emergence of scale invariance and long-range correlations (Supplementary Fig. 2 and 3). Screening of subject-specific parameter spaces showed that the power-law exponent of simulated fMRI depended on the balance of large-scale excitation and local inhibition: the 2D distribution of prediction quality (fMRI time series and functional connectivity) and critical exponent showed a characteristic diagonal pattern, which demonstrates the necessity of proper balance of large-scale excitation and local inhibition for optimal prediction (Supplementary Fig. 3).

**Figure 6.**
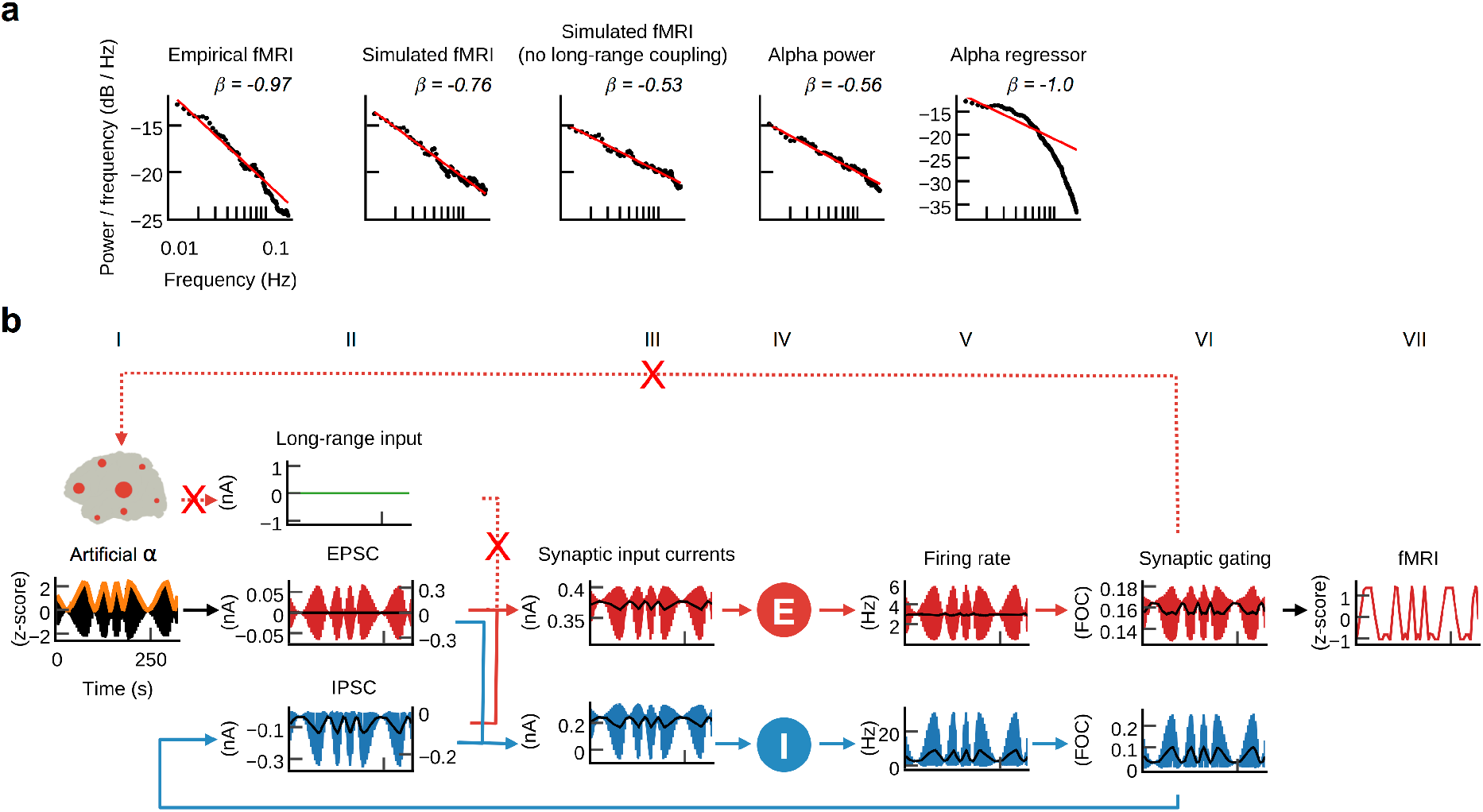
Large-scale coupling controls scale-free fMRI dynamics. **a**, Empirical fMRI power spectra show power-law scaling with exponent *β_emp_* = −0.97. In accordance, the power-law exponent of simulated fMRI was close to the empirical value (*β_sim_* = −0.76). Previous results showed that the model transformed ongoing EEG source power fluctuation into fMRI oscillations. Notably, however, alpha power time series had a considerably flatter power spectrum slope (*β_α-band_* = −0.56, *β_wide-band_* = −0.53) than fMRI. Simulation results that were obtained for deactivated large-scale coupling had a similarly flat spectrum (*β_zeroG_* = −0.53). Parameter space exploration suggests that the emergence of scale-free fMRI power spectra depends on the proper balancing of recurrent large-scale excitation with local inhibition (Supplementary Figs. 2 and 3). When large-scale coupling was absent or inadequately balanced, prediction quality decreased and models produced flatter spectra or no scale invariance at all. **b**, Grand average waveforms of simulation results using the same artificial α-rhythm input as in Fig. 5b, but with disabled large-scale coupling. In contrast to Fig. 5b, the resulting firing rates, synaptic gating and fMRI waveforms showed no frequency-amplitude dependence. Comparison of model dynamics between both scenarios suggests that gradually accumulating self-reinforcing excitation through large-scale coupling leads to frequency-dependent amplification that augments slower oscillations more than faster oscillations, which results in the emergence of scale invariance.

## DISCUSSION

Connectome-based computational modelling has helped to provide new insights into large-scale network dynamics related to phenomena such as resting-state networks^45^. We build from this foundation by more explicitly integrating empirical data to constrain the model dynamics. Subject-specific connectomes and biophysically based brain models were driven by their own EEG-derived approximations of locally generated synaptic input currents and fit to their own fMRI time series. The optimized models enabled inferences about mesoscopic dynamics underlying fMRI resting state activity. We reinforce the validity of the model by showing that the underlying emerging processes, which were not explicitly incorporated in the model, reproduce neurophysiological results. The novel approach forms an integrative framework to construct hypotheses derived from the generated model activity, while remaining sensitive to constraints provided by empirical evidence. Thereby, it links data to theory by uniting models of neural dynamics with empirical observations across modalities and spatiotemporal scales.

We demonstrate how this framework can be used for the systematic study of individual brain activity by identifying neural mechanisms underlying person-specific resting-state fMRI activity, the inhibitory effect of EEG α-rhythms and the emergence of scale-free dynamics. Power-law relationships between frequency and amplitude of oscillations are omnipresent in nature; the identified mechanism—increased amplification of slower processes through prolonged recurrent excitation—may have implications for other areas of science. The observed co-emergence of long-range correlations and power-law scaling may point to a unifying explanation within the theory of self-organized criticality, as previously proposed by others^46^.

Fitting models with subject-specific fMRI time series configured network interaction such that mesoscopic dynamics emerged that are consistent with other empirical data at that scale. Estimated parameters, like the strengths of local and global coupling, created dynamical regimes that were characterized by ongoing E/I balance, which we also found to be a key mechanism underlying fMRI resting-state oscillations and α-inhibition. Our findings mechanistically link electrical activity to hemodynamic oscillations and thereby add to accumulating evidence suggesting that RSNs originate from neuronal activity^11–13, 25, 26, 29, 30^ rather than being a purely hemodynamic phenomenon that is only correlated, but not caused by it^32, 47, 48^. Likewise, our results provide a mechanistic explanation for the emergence of power-law spectra that is based on large-scale interaction of brain regions. Our identified mechanism suggests a neural origin in line with empirical results that connect E/I balance to scale-free dynamics^49^ and recurrent excitation to states of high or low activity in local circuits^50^. Our major results are integrated and visualized in **Supplementary Video 1**.

Source activity injection aims to increase the biological realism of brain simulation. Computational models by necessity omit features of the real system for the sake of simplicity, generality and efficiency. Adding such features increases degrees of freedom, but may also render parameter spaces intractable and increases the risk of over-fitting. Injection of empirical activity is a way to systematically probe sufficiently abstract neural systems while maintaining biologically realistic behaviour. Thereby, the approach aims to balance a level of abstraction that is sufficient to provide insights with being detailed enough to guide subsequent empirical study.

The present approach proved promising for general decoding of mesoscopic neural information processing mechanisms. The ability to selectively target and perturb person-specific whole-brain models will contribute to our mechanistic understanding of neural information processing and its relation to brain function and dysfunction.

## METHODS

### Computational model

The model used in this study is based on the large-scale dynamical mean field model used by Deco and colleagues^51, 52^. Brain activity is modelled as the network interaction of local population models that represent cortical areas. Cortical regions are modelled by interconnected excitatory and inhibitory neural mass models. In contrast to the original model, excitatory connections were replaced by injected EEG source activity. The dynamic mean field model faithfully approximates the time evolution of average synaptic activities and firing rates of a network of spiking neurons by a system of coupled non-linear differential equations for each node *i*:

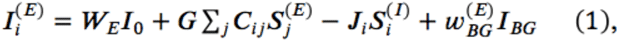

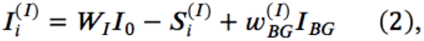

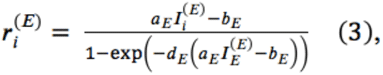

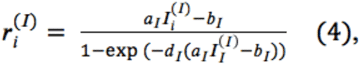

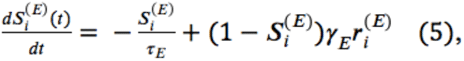

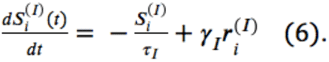

Here, 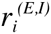 denotes the population firing rate of the excitatory *(E)* and inhibitory *(I)* population of brain area *i*. 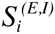 identifies the average excitatory or inhibitory synaptic gating variables of each brain area, while their input currents are given by 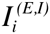. In contrast to the model used by Deco et al.^51^ that has recurrent and feedforward excitatory coupling, we approximate excitatory postsynaptic currents *I_BG_* using region-wise aggregated EEG source activity that is added to the sum of input currents 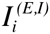. This approach is based on intracortical recordings that suggest that EPSCs are non-random, but strongly correlated with electric fields in their vicinity, while IPSCs are anticorrelated with EPSCs^8–10^. The weight parameters 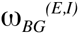 rescale the z-score normalized EEG source activity independently for excitatory and inhibitory populations. *G* denotes the large-scale coupling strength scaling factor that rescales the structural connectivity matrix *C_ij_* that denotes the strength of interaction for each region pair and *j*. All three scaling parameters are estimated by fitting simulation results to empirical fMRI data by exhaustive search. Initially, parameter space (*n*-dimensional real space with *n* being the number of optimized parameters) was constrained such that the strength of inhibition was larger than the strength of excitation, satisfying a biological constraint. Furthermore, for each tested parameter set (containing the three scaling parameters mentioned above) the region-wise parameters *J_i_* that describe the strength of the local feedback inhibitory synaptic coupling for each area (expressed in nA) are fitted with the algorithm described below such that the average firing rate of each excitatory population in the model was close to 3.06 Hz (i.e. the cost function for tuning parameters *J_i_* was solely based on average firing rates and not on prediction quality). The overall effective external input *I_0_*=0.382 nA is scaled by *W_E_* and *W_I_*, for the excitatory and inhibitory pools, respectively. 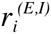 denotes the neuronal input-output functions (f-I curves) of the excitatory and inhibitory pools, respectively. All parameters except those that are tuned during parameter estimation are set as in Deco et al.^51^. BOLD activity was simulated on the basis of the excitatory synaptic activity *S^(E)^* using the Balloon-Windkessel hemodynamic model^41, 53^.

### Feedback inhibition control

Using standard parameters of the original model, the excitatory populations of isolated nodes have an average firing rate of 3.06 Hz, which conforms to the Poisson-like cortical *in vivo* activity of ∼3 Hz^54–56^. For coupled populations, firing rates change in dependence of the employed structural connectivity matrix and injected input. To compensate for a resulting excess or lack of excitation, a local regulation mechanism, called feedback inhibition control (FIC), was used. The approach was previously successfully used to significantly improve FC prediction as well as for increasing the dynamical repertoire of evoked activity and the accuracy of external stimulus encoding^51^. Despite the mentioned advantages of FIC tuning, it brings the disadvantage of increasing the number of open parameters of the model. To prove that prediction quality is not due to FIC, but solely due to the three global parameters and to exclude concerns about over-parameterization or that FIC may be a potentially necessary condition for the emergence of scale-freeness, we devised a control model that did not implement FIC, but used a single global parameter for inhibitory coupling strength. Instead of tuning the 68 individual local coupling weights individually, only a single global value for all inhibitory coupling weights *J_i_* was varied. We compared the effect of FIC on time series prediction quality and found no significant difference in prediction quality to simulations that used only a single value for all local coupling weights *J_i_* per subject (*p = .92, two-sample t-test*). In contrast to simulations that are driven by noise^51^, FIC parameters for injected input must be estimated for the entire simulated time series, since the non-stationarity of stimulation time series leads to considerable fluctuations of firing rates.

Therefore, we developed a local greedy search algorithm for fast FIC parameter estimation based on the algorithm in Deco et al.^51^. To exert FIC, local inhibitory synaptic strength is iteratively adjusted until all excitatory populations attained a firing rate close to the desired mean firing rates for the entire ∼20 minutes of activity. During each iteration, the algorithm performs a simulation of the entire time series. Then, it computes the mean firing activity over the entire time series for each excitatory population and adapts *J_i_* values accordingly, i.e., it increases local *J_i_* values if the average firing rate over all excitatory populations during the *k*-th iteration 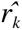 is larger than 3.06 Hz and vice versa. In contrast to the algorithm by Deco et al., the value by which *J_i_* is changed is dynamically adapted in dependence of the firing rate obtained during the current iteration

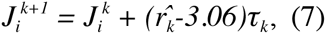
where 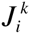 denotes the value of feedback inhibition strength of node *i* and *τ_k_* denotes the adaptive tuning factor during the *k*-th iteration. In the first iteration, all *J_i_* values are initialized with 1 and *τ_k_* is initialized with 0.005. The adaptive tuning factor is dynamically changed during each iteration based on the result of the previous iteration:

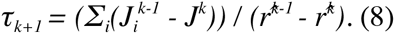

For the case that the result did not improve during the current iteration, i.e.,

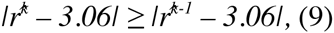
the adaptive tuning factor is decreased by multiplying it with 0.5 and the algorithm continues with the next iteration. After 12 iterations, all *J_i_* values are set to the values they had during the iteration *k* where *|r^k^ − 3.06|* was minimal.

### MRI preprocessing

Structural and functional connectomes from 15 healthy subjects (age range: 18 – 31 y., 8 female) were extracted from full data sets (diffusion-weighted MRI, T1-weighted MRI, EEG-fMRI) using a local installation of the automatic pipeline developed by Schirner et al.^57^. MRI measurements were acquired with 32-channel Siemens 3 Tesla Trio systems at the Berlin Center for Advanced Neuroimaging, Berlin, Germany. Extracted structural connectivity approximates the strengths and time-delays of interaction between regions as mediated by white matter fiber tracts. Heterogeneous conduction delays between distant cortical areas are neglected in this study and set to 1 ms for each connection. Strength matrices *C_ij_* were divided by their respective maximum value for normalization. In short, the pipeline proceeds as follows: for each subject a three-dimensional high-resolution T1-weighted image (1 mm isotropic) image was used to divide cortical gray matter into 68 regions according to the Desikan-Killiany atlas using FreeSurfer’s^58^ automatic anatomical segmentation and registered to diffusion data. The gyral-based brain parcellation is generated by an automated probabilistic labelling algorithm that has been shown to achieve a high level of anatomical accuracy for identification of regions while accounting for a wide range of inter-subject anatomical variability^59^. The atlas was successfully used in previous modelling studies and provided highly significant structure-function relationships^4, 57, 60^. Probabilistic white matter tractography and track aggregation between each region-pair was performed as implemented in the automatic pipeline and the implemented distinct connection metric extracted. This metric weights the connection strength between two regions according to the gray-matter/white-matter interface areas of both regions used to connect these regions and not by the number of raw tracks, since this number is biased by different anatomical features^57^. After preprocessing, the cortical parcellation mask was registered to fMRI resting-state data (T2-weighted echo planar imaging with blood oxygen level-dependent contrast; TR = 1940 ms; voxel size = 3 mm isotropic; eyes-closed resting-state) of subjects and average fMRI signals for each region were extracted. The first five images of each scanning run were discarded to allow the MRI signal to reach steady state. To identify RSN activity a spatial Group ICA decomposition was performed for the fMRI data of all subjects using FSL MELODIC (MELODIC v4.0; FMRIB Oxford University, UK, Beckmann and Smith (61)) with the following parameters: high pass filter cut off: 100 s, MCFLIRT motion correction, BET brain extraction, spatial smoothing 5 mm FWHM, normalization to MNI152, temporal concatenation, dimensionality restriction to 30 output components. ICs that correspond to RSNs were automatically identified by spatial correlation with the nine out of the ten well-matched pairs of networks of the 29,671-subject BrainMap activation database as described in Smith et al.^62^ (excluding the cerebellum network).

### EEG preprocessing

Details of EEG preprocessing are described in supplementary material of Schirner et al.^57^. First, to account for slow drifts in EEG channels all channels were high-pass filtered at 1.0 Hz (standard FIR filter). Imaging Acquisition Artefact (IAA) correction was performed using Analyser 2.0 (v2.0.2.5859, Brain Products, Gilching, Germany). The onset of each MRI scan interval was detected using a gradient trigger level of 300 μV/ms. Incorrectly detected markers, e.g. due to shimming events or heavy movement, were manually rejected. To assure the correct detection of the resulting scan start markers each inter-scan interval was controlled for its precise length of 1940 ms (TR). For each channel a template of the IAA was computed using a sliding average approach (window length: 11 intervals) and subsequently subtracted from each scan interval. For further processing, the data was down sampled to 200 Hz, imported to EEGLAB and low-pass filtered at 60 Hz. ECG traces were used to detect and mark each instance of the QRS complex in order to identify ballistocardiogram (BCG) artifacts. The reasonable position and spacing of those ECG markers was controlled by visual inspection and corrected if necessary. To correct for BCG and artifacts induced by muscle activity, especially movement of the eyes, a temporal ICA was computed using the extended Infomax algorithm as implemented in EEGLAB. To identify independent components (ICs) that contain BCG artifacts the topography plot, activation time series, power spectra and heartbeat triggered average potentials of the resulting ICs were used as indication. Based on established characteristics, all components representing the BCG were identified and rejected, i.e., the components were excluded from back-projection. The remaining artificial, non-BCG components, accounting for primarily movement events especially eye movement, were identified by their localization, activation, spectral properties and ERPs. Detailed descriptions of EEG and fMRI preprocessing have been published elsewhere^27, 63–65^.

### Biologically based model input

EEG source imaging was performed with the freely available MATLAB toolbox Brainstorm using default settings and standard procedure for resting-state EEG data as described in the software documentation^66^. Source space models were based on the individual cortical mesh triangulations as extracted by FreeSurfer from each subject’s T1-weighted MRI data and downsampled by Brainstorm. From the same MRI data, head surface triangulations were computed by Brainstorm. Standard positions of the used EEG caps (Easy-cap; 64 channels, MR compatible) were aligned by the fiducial points used in Brainstorm and projected onto the nearest point of the head surface. Forward models are based on Boundary Element Method head models computed using the open-source software OpenMEEG and 15002 perpendicular dipole generator models located at the vertices of the cortical surface triangulation^67, 68^. The sLORETA inverse solution was used to estimate the distributed neuronal current density underlying the measured sensor data since it has zero localization error^69^. EEG data was low-pass filtered at 30 Hz and imported into Brainstorm. There, the epochs before the first and after the last fMRI scan were discarded and the EEG signal was time-locked to fMRI scan start markers. Using brainstorm routines, EEG data was projected onto the cortical surface using the obtained inversion kernel and averaged according to the Desikan-Killiany parcellation that was also used for the extraction of structural and functional connectomes and region-averaged fMRI signals. The resulting 68 region-wise source time series were imported to MATLAB, z-score normalized and upsampled to 1000 Hz using spline interpolation as implemented by the Octave function *interp1*. To enable efficient simulations, the sampling rate of the injected activity was ten times lower than model sampling rate. Hence, during simulation identical values have been injected during each sequence of ten integration steps.

### Simulation and analysis

Simulations were performed with a highly optimized C implementation of the used model on the supercomputers JUROPA and JURECA at the Juelich Supercomputing Centre. The model was implemented using several code optimization strategies that considerably increase execution speed, which enabled an exhaustive brute-force parameter space scan using 3888 combinations of the parameters *G* and 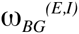 for each subject. Each of these combinations was computed 12 times to iteratively tune *J_i_* values. As control setup, further simulations were performed with random permutations of the input time series. Therefore, each source activity time series was randomly permutated (individually for each region and subject) using the Octave function *randperm()* and injected into simulations using all parameter combinations that were previously used. As an additional control situation the original dynamic mean field model as described in Deco et al.^51^ was simulated for the 15 SCs. Here, the parameters *G* and *J_NMDA_* were varied and FIC tuning was performed using the same algorithm as used for the source activity injection model. The simulation and FIC optimization process was identical for all three models. The length of the simulated time series for each subject was 21.6 minutes. Simulations were performed at a model sampling rate of 10,000 Hz. BOLD time series were computed for every 10^th^ time step of excitatory synaptic gating activity using the Balloon-Windkessel Model^41^. From the resulting time series every 1940^th^ step was stored in order to obtain a sampling rate of simulated fMRI that conforms to the empirical fMRI TR of 1.94 s. The first 11 scans (21.34 s) of activity were discarded to allow model activity and simulated fMRI signal to stabilize. For each subject and modelling approach the simulation result that yielded the highest average correlation between all 68 empirical and simulated regions time series for all tested parameters was used for all analyses. A five-fold cross-validation scheme was performed on the EEG injection model data. Therefore, the data was divided into two subsets: 80 % as training subset and 20 % as testing subset. Prediction quality was estimated using the training set, before trained models were asked to predict the testing set. Resulting prediction quality was compared between training and test data set and between test data set and the data obtained from fitting the full time series. To assess time-varying prediction quality, a correlation analysis was performed in which a window with a length of 100 scans (194 s) was slid over the 68 pairs of empirical and simulated time series and the average correlation over all 68 regions was computed for each window. For RSN analysis, only those regions were compared with the temporal modes of RSNs that had a spatial overlap of at least 40 % of all voxels belonging to the respective region. For the estimation of signal correlation, the computation of entries of FC matrices and as a measure of similarity of FC matrices Pearson’s linear correlation coefficient was used. FC matrices were compared by stacking all elements below the main diagonal into vectors and computing the correlation coefficient of these vectors. FC dynamics prediction quality was estimated by computing the mean correlation obtained for all window-wise correlations of a sliding window analysis of empirical and simulated time series (window-size: 100 scans = 194 s).

For computation of average power spectral densities (PSDs) of empirical and simulated signals, region time series were selected using rigorous model selection criteria to ensure scale invariance of the analysed time series; on average 79 % of all 1020 region-wise time series (15 subjects x 68 regions) for the seven analysed signal types (empirical fMRI, simulated fMRI, simulated fMRI without global coupling, simulated fMRI without FIC, simulated fMRI without FIC and without global coupling, alpha power, alpha-regressor) tested as scale-free; for every signal type every subject had at least five regions to test as scale-free. PSDs were computed using the Welch method as implemented in Octave, normalized by their total power and averaged. Resulting average frequency spectra were fitted with a power-law function *f(x) = ax^β^* using least-squares estimation in the frequency range 0.01 Hz and 0.17 Hz which is identical to the range for which the test for scale invariance was performed. Frequencies below were excluded in order to reduce the impact of low-frequency signal confounds and scanner drift, frequencies above that limit were excluded to avoid aliasing artefacts in higher frequency ranges (TR = 1.94 s, hence Nyquist frequency is around 0.25 Hz). In order to compare the scale invariance of our empirical fMRI data with results from previous publications^44^, we also computed spectra in a range that only included frequencies < 0.1 Hz.

In order to adequately quantify scale invariance we applied rigorous model selection to every time series to identify power-law scaling and excluded all time series from analyses that were described better by a model other than a power-law. Nevertheless, we compared the obtained results from this strict regime with results obtained when all time series were included and found them to be qualitatively identical. To test for the existence of scale invariance we used a method that combines a modified version of the well-established detrended fluctuation analysis (DFA) with Bayesian model comparison^70^. DFA is, in contrast to PSD analyses, robust to both stationary and nonstationary data in the presence of confounding (weakly non-linear) trends. Rather than averaging the mean squared fluctuations over consecutive intervals as in conventional DFA, this method uses the values per interval to approximate the distribution of mean squared fluctuations with kernel density estimation. This allows for estimating the corresponding log-likelihood as a function of interval size without presuming the fluctuations to be normally distributed, as in the case of conventional DFA, which gives a non-parametric estimate of the log-likelihood for fitted models. Furthermore, conventional DFA does not provide any means to determine whether a power law is present or not. It is important to note, that a simple linear fit of the detrended fluctuation curve without proper comparison of the obtained goodness of fit with that of other models would entirely ignore alternative representations of the data different than a power law. For quantification of the goodness of fit with simple regression its corresponding coefficient of determination, *R^2^*, is ill-suited as it measures only the strength of a linear relationship and is inadequate for nonlinear regression^71^. Here, we assess power-law scaling in the context of DFA, i.e. the optimality of a straight line fit of fluctuation magnitude against interval size in a log-log representation, with non-parametric model selection using the Bayesian information criterion in order to compare the linear model against alternative models. Details of the used method can be found elsewhere^70^. Briefly, the method first estimates the cumulative sum of each time series. Next, signals are divided into non-overlapping intervals of increasing length, for a range of window sizes (48 time windows in steps from 3 to 50 data points). Then, for each interval the linear trend is removed and the root mean squared (RMS) fluctuation for each detrended interval is computed. Interval size is plotted against the RMS magnitude of fluctuations in a log-log representation and model-fits with 11 different models are computed using maximum likelihood estimation and compared to the linear fit. A straight line on the log-log graph indicates scale invariance expressed as *F(n)αn^α^*, with *n* the interval size, *F(n)* detrended fluctuation and α, the scaling exponent, which represents the slope of the straight line fit (*α ≅ 0.5*, indicates uncorrelated white noise, while in the case of *α > 0.5* the auto-correlation function decays slower than the auto-correlation function of Brownian motion, indicating long-term ‘memory’). Lastly, likelihoods are used to compute Bayesian information criteria (BIC) for each model, which are used to select among models. BIC take into account model accuracy (as quantified by maximum likelihood) and model complexity, which scores the number of free parameters used in the different models. Optimality in the context of BIC therefore yields a maximally accurate while minimally complex explanation for data, i.e., the optimal compromise between goodness-of-fit and parsimony. For the different signals the majority of time series were tested as being scale free: 83 % for empirical fMRI, 69 % for simulated fMRI, 71 % for simulated fMRI with deactivated FIC, 83 % for simulated fMRI with deactivated global coupling, 86 % for simulated fMRI with deactivated global coupling and FIC, 90 % for alpha power and 70 % for the alpha-regressor.

To compute grand average waveforms, state-variables were averaged over all 15 subjects and 68 regions (N=1020 region time series) time-locked to the zero crossing of the α-amplitude, which was obtained by band-pass filtering source activity time series between 8 and 12 Hz; to obtain sharp average waveforms, all α-cycle epochs with a cycle length between 95 and 105 ms were used (N=4,137,994 α-cycle). For computing ongoing α-power time courses, instantaneous power time series were computed by taking the absolute value of the analytical signal (obtained by the Hilbert transform) of band-pass filtered source activity in the 8 – 10 Hz frequency range; the first and last ∼50 s were discarded to control for edge effects. To compute the alpha regressor, power time series were convolved with the canonical hemodynamic response function, downsampled to fMRI sampling rate and shifted relative to fMRI time series to account for the lag of hemodynamic response. The highest negative average correlation over all 68 region-pairs obtained within a range of +/−3 scans shift was used for comparison with simulation results.

---

**Supplementary Figure 1.**
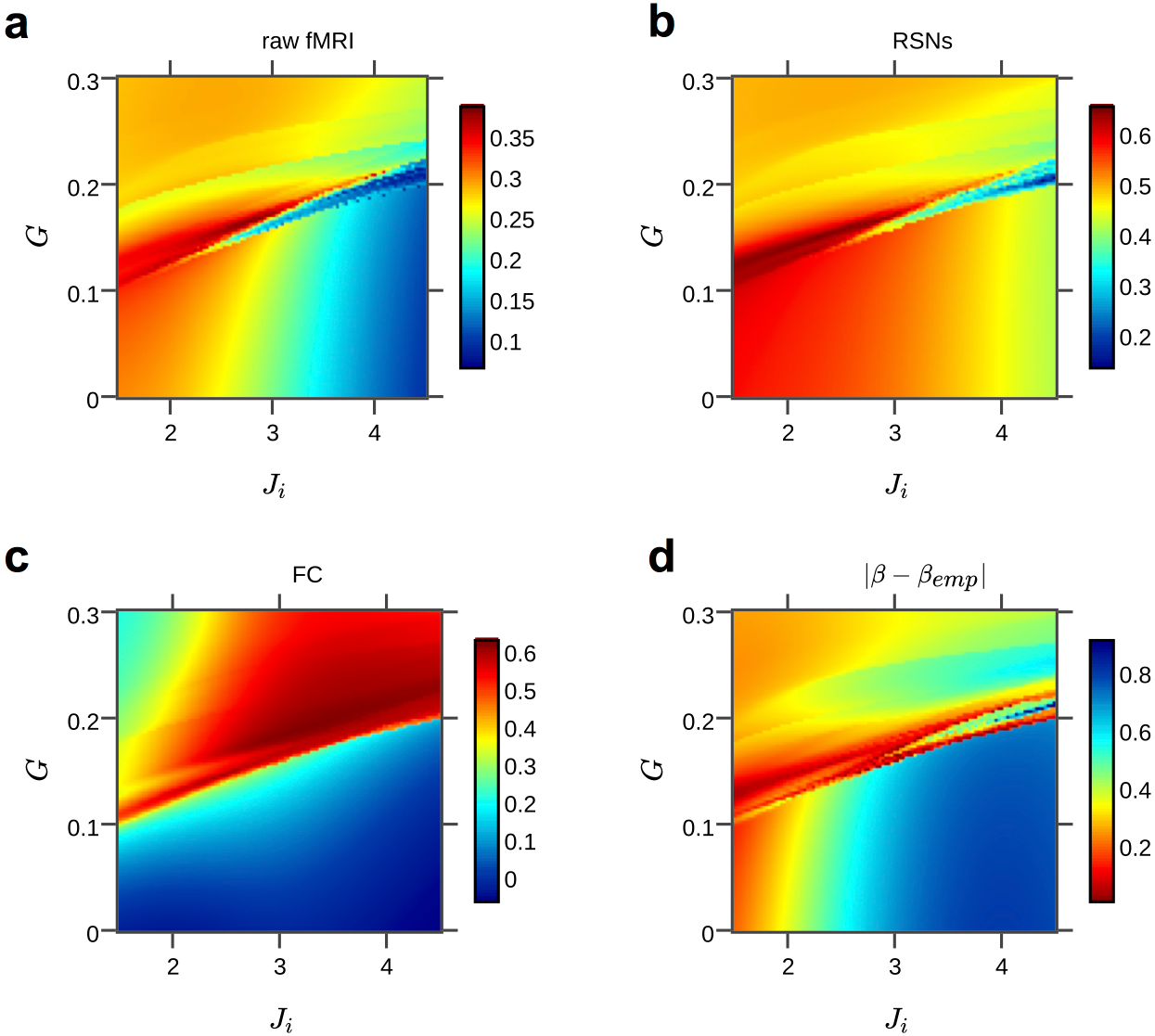
Parameter space exploration results. **a-c**, 2d parameter space heat maps show average time series correlation over all 68 regions obtained from the hybrid model for different combinations of the three varied parameters *G*, 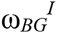 and 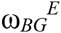 (the latter depicted as ratio 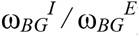); results were averaged over all subjects. Parameter values that yielded the highest average correlation were used for simulations with artificial alpha input (marked with an asterisk). We confirmed identifiability of the model by showing that parameter space search converges towards a single optimal solution yielding best predictions.

**Supplementary Figure 2.**
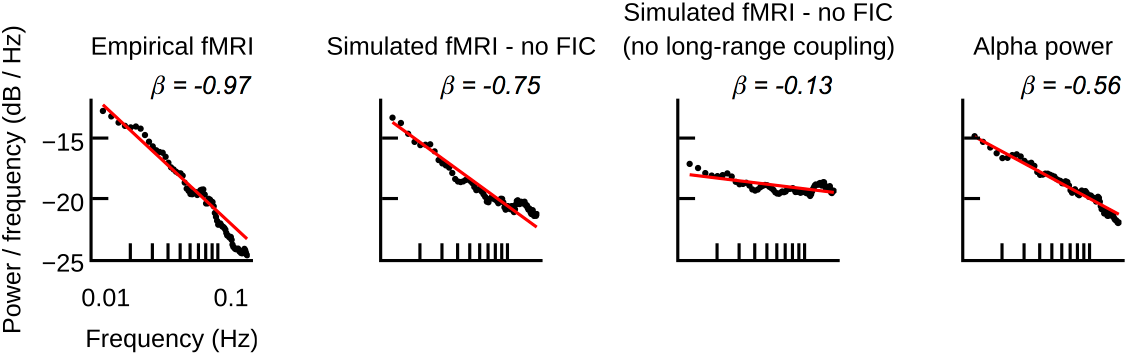
Power spectral densities (averaged over all subjects and regions) and power-law fits for empirical and simulated fMRI, ongoing alpha power time series of EEG source activity input and the alpha regressor. For comparison with simulation results shown in Figure 6a, the models used here implemented no feedback inhibition control (FIC), i.e., a single value for all local inhibitory connection parameters *J_i_* was used. Empirical and simulated fMRI spectra have a large power-law exponent, i.e. a steeper slope, compared to ongoing alpha power or wide-band power (*β_α-band_* = −0.56, *β_wide-band_* = −0.53). To analyse the effect of large-scale network interaction, simulated fMRI was computed with and without long-range coupling. To exclude that power-law spectra emerge despite absent large-scale coupling, local inhibition was tuned such that the model produced highest fMRI time series predictions. Without long-range coupling, simulated fMRI showed a similar exponent as injected source activity. When large-scale coupling was activated and global excitation was properly balanced with local inhibition, the exponent was closer to the exponent of empirical fMRI (Suppl. Fig. 3).

**Supplementary Figure 3.**
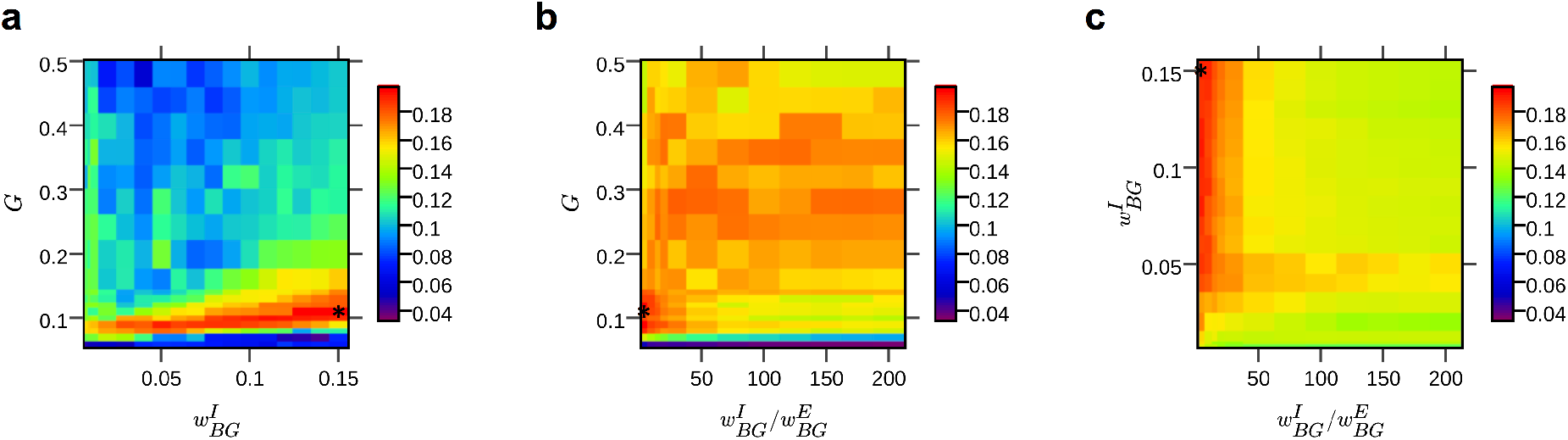
Parameter space exploration of the simplified hybrid model (without FIC, i.e., using a single value for all local inhibitory connection parameters *J_i_*) for an exemplary subject. Colours indicate the average correlations between simulated and empirical data (**a-c**) and the absolute difference between exponents of power-law fits with empirical and simulated power spectral densities (**d)**, averaged over all regions. The distributions of the different metrics suggest a link between prediction quality (of raw fMRI, RSNs and FC and the power-law exponent *β*) and the relative strengths of long-range coupling *G* and local inhibition *J_i_*. A diagonal pattern in heat maps indicates that prediction quality and power-law exponents depend on the balancing of large-scale excitation with local inhibition. The plots illustrate that when large-scale coupling was absent or inadequately balanced, models did not produce scale-free behaviour and prediction quality decreased.

## Acknowledgments

Computations were performed on JUROPA and JURECA at Forschungszentrum Juelich (www.fz-juelich.de, Grant NIC#8344 & NIC#10276 to P.R.). The authors acknowledge the support of the James S. McDonnell Foundation (Brain Network Recovery Group JSMF22002082) to A.R.M., V.J., G.D., and P.R., the German Ministry of Education and Research (US-German Collaboration in Computational Neuroscience to and Bernstein Focus State Dependencies of Learning 01GQ0971-5, the Max-Planck Society and funding from the European Union Horizon2020 (ERC Consolidator grant) to P.R. The authors thank Olaf Sporns, Jochen Braun and Andreas Daffertshofer for their helpful comments on the manuscript.

## Author contributions

M.S. and P.R. conceived the project. M.S. conceived and implemented the model based on models and code by D.G., V.J. and A.R.M. M.S. assembled input data from previously collected empirical data (Schirner et al. 2015). P.R. supervised the project. M.S. and P.R. analysed and interpreted the results. M.S. wrote the paper with contributions from P.R., A.R.M., V.J. and G.D. All authors discussed the results and commented on the manuscript at all stages.

## Author information

The authors declare no competing financial interests. Correspondence and request for materials should be addressed to P.R. or M.S.

